# Interaction Effects of Apolipoprotein E *ε* 4 and Cognitive Status on the Functional Connectivity within Default Mode Network in Individuals at Risk of Cognitive Impairments

**DOI:** 10.1101/565812

**Authors:** Hanna Lu, Suk Ling Ma, Winnie Chu Chiu Wing, Savio Wai Ho Wong, Linda C. W. Lam

## Abstract

**Background:** Disturbance of intrinsic brain networks is often associated with APOE *ε* 4 allele and cognitive dysfunction. However, little is known about the functional connectivity strength (FCS) within default mode network (DMN).

**Objective:** We aimed to examine the independent effects APOE *ε* 4 and cognitive status and the interaction effect on the functional connectivity within DMN.

**Methods:** Resting-state functional MRI was conducted for sixty-five senior adults who had normal cognition or cognitive decline with or without APOE *ε* 4. Regions within DMN include mPFC, IPL, LTC, hippocampus and PCC. The absolute values of correlation coefficient between DMN regions were employed as the measures of FCS for quantifying the functional connectivity.

**Results:** Main effect of APOE *ε* 4 was found on the FCS of bilateral PCC (*F* = 6.133, *p* = 0.016), while the main effect of cognitive status was found on the FCS of left IPL and right mPFC (*F* = 4.585, *p* = 0.036). Interaction effect was found in the FCS of right mPFC and left LTC (*F* = 4.698, *p* = 0.034), right hippocampus and left LTC (*F* = 7.673, *p* = 0.008), left PCC and left LTC (*F* = 6.719, *p* = 0.012), right IPL and right LTC (*F* = 4.516, *p* = 0.038).

**Conclusions:** APOE *ε* 4 carriers with cognitive impairment denote a preclinical status characterized by aberrant inter-hemispheric FC within DMN. The network-level connectivity may be useful in the evaluation of the individuals at risk for developing AD and affiliate network-guided brain stimulation.

## INTRODUCTION

Research into the presymptomatic stages of Alzheimer’s disease (AD) and the discovery of status-specific biomarkers is of utmost importance to make a more accurate diagnosis at the very onset of the disease. With the advances of neuroimaging techniques, AD has been reframed as a disease of ‘disconnectivity’ with the presence of abnormal functional connectivity [1,2]. Default mode network (DMN), as a core intrinsic brain network, is vulnerable to aging and neuropathological cascade [3]. Compelling evidence illustrates that beta-amyloid (A*β*), as a pathological hallmark of AD, deposits and spreads progressively within DMN across the lifespan [4].

The Apolipoprotein E (APOE) *ε* 4 allele is one of the well-replicated genetic risk factor for sporadic AD [5,6], attending the formulation of A*β* [7]. As to the effect of APOE *ε* 4 on DMN, both reduced and increased functional connectivity (FC) were observed in cognitively normal APOE *ε* 4 carriers compared with their peers [8–11]; while increased FC was only found in young carriers [12,13]. These results indicate that there might be a status-dependent compensatory mechanism to maintain brain function decades before showing clinical manifestation.

Indeed, the linkage between APOE *ε* 4 and DMN connectivity becomes more complicated in late adulthood. A promising avenue to better understand this complex linkage is to examine the changes from two levels: (1) Status-level: Cross-sectional studies reported decreased FC within DMN over time [14–17]. Recently, a longitudinal study also confirmed the reduced connectivity within DMN in an ageing cohort [4]. (2) Network-level: Stemming from the heterogeneity within DMN, two sub-networks, including anterior DMN (aDMN) and posterior DMN (pDMN), has been highlighted regarding to their distinct connection patterns [18,19]. Several studies found AD-specific disturbance of FC within DMN [20,21].

Accordingly, it should be noted that decreased DMN connectivity is not restricted to APOE *ε* 4, but also interferes with cognitive status. Therefore, the pending question would be whether APOE *ε* 4 carriers with or without cognitive deficits display the status-specific FC within DMN, and to what extent the FC correlate with cognitive functions? In this study, we sought to examine the alternations of functional connectivity strength (FCS) of DMN connectivity in senior APOE *ε* 4 carriers with or without cognitive decline and evaluate the effect of interaction between APOE *ε* 4 and cognitive status on DMN connectivity. We hypothesized that APOE *ε* 4 MCI carriers would show reduced FCS within DMN. Moreover, we hypothesized that the effect of ‘APOE *ε* 4 × status’ interaction on FSC of DMN connectivity would be associated with impaired cognitive function.

## Materials and Methods

### Participants

Sixty five right-handed Chinese senior adults (aged from 66 to 82 years) were recruited our cohort [22], including 38 cognitively normal (CN) adults and 27 mild cognitive impairment (MCI). CN adults were defined by a score of Mini Mental State Examination (MMSE) greater than 28. MCI patients were determined by the following three items [23]: (1) With the score of cognitive performance within 1.5 standard deviation of age and education matched normative values derived from the cohort [22]; (2) No interference with independence in daily activities and (3) No better explanation by other mental disorders. The cases showing depressive disorder, sleep problems or a history of neurological or psychiatric disorders were excluded.

All participants underwent a comprehensive battery of neuropsychological tests, APOE genotyping and magnetic resonance imaging (MRI) assessments including structural MRI (sMRI) and resting-state functional MRI (rs-fMRI). All participants have signed the written informed consent before APOE genotyping and MRI assessments.

Ethics approval was obtained from the Joint Chinese University of Hong Kong-New territories East Cluster Clinical Research Ethics Committee (Joint CUHK-NTECCREC).

### Evaluation of cognitive performance

A comprehensive battery of neuropsychological tests was conducted to assess the main domains of cognitive functions [17]. Alzheimer’s Disease Assessment Scale cognitive subscale (ADAS-Cog) and Mini mental state examination (MMSE) were used to measure the global cognitive efficiency. Delayed recall of words and digit span backward (DSB) were used to evaluate working memory function. Trail making test part A (TMT-A) and digit span forward (DSF) were employed to assess attention. Trail making test part B (TMT-B) and category verbal fluency test (CVFT) were used to assess executive function. All the measurements were conducted with Chinese instructions.

### APOE genotyping

All participants provided a sample of DNA for APOE genotyping. Genomic DNA was extracted from peripheral blood samples using DNA extraction kit according to the instruction (Qiagen, US). The presence of APOE *ε* 4 allele was analysed and performed using the well-validated PCR-RFLP method [24].

### Image acquisition

All MRI data were collected in the Prince of Walsh Hospital by using a 3 Tesla Philips Achieva Scanner with an eight-channel SENSE head coil. A series of T1-weighted structural images were obtained for each participant with the following parameters: Repetition Time (TR) = 7.4 ms, Echo Time (TE) = 3.4 ms, flip angle = 8°, voxel size = 1.04 × 1.04 × 0.6 mm^3^.

During the session of rs-fMRI, all participants were instructed to relax and keep their eyes close and would immediately addressed after each rs-fMRI run via the intercom to ensure they stay awake in MRI scanner. In each scanning sequence, we obtained a series of 40 axial T2-weighted gradient echo-planar images (TR = 3000 ms, TE = 25 ms, flip angle = 90°, field of view (FOV) = 230 × 230 mm^2^, in-plane resolution = 2.4 × 2.4 mm^2^, matrix = 96 × 96, slice thickness = 3 mm). A total of 128 time points of rs-fMRI were acquired for each participant.

### Voxel-based morphometry (VBM) analysis

Structural MRI (sMRI) data were pre-processed by VBM using Statistical Parametric Mapping (SPM12) (Wellcome Department of Imaging Neuroscience, London, United Kingdom). All sMRI images were reoriented by anterior commissure - posterior commissure line (AC-PC line) manually and segmented into grey matter (GM), white matter (WM), and cerebrospinal fluid. GM images were then normalized to standard Montreal Neurological Institute (MNI) space using the Diffeomorphic Anatomical Registration Through Exponentiated Lie Algebra (DARTEL) [25]. Voxel volumes were modulated by whole brain volume with proportional scaling. All images were smoothed using a Gaussian kernel of 8 × 8 × 8 mm full width at half maximum (FWHM). VBM analyzed differences in local GM and WM volumes across the whole brain using analysis of variance (ANOVA) across all four groups.

### Functional connectivity analysis

#### Pre-processing

Pre-processing of rs-fMRI data was conducted by the Statistical Parametric Mapping software (SPM12, http://www.fil.ion.ucl.ac.uk/spm/software/spm12/) and Resting-State fMRI Data Analysis Toolkit plus V1.21 (REST plus V1.21, http://restfmri.net/forum/RESTplusV1.2) [26] embedded in MATLAB R2016a. The first 5 volumes of each functional time points were discarded because of the instability of the initial MRI signal and initial adaption of participants to the situation. The remaining rs-MRI data was subsequently corrected for slice timing and realigned to the first image by rigid-body head movement correction. Then, all rs-fMRI images were coregister to high-resolution T1 images and normalized to standard stereotaxic anatomical Montreal Neurological Institute (MNI) space. The normalized volumes were spatially smoothed using an isotropic Gaussian filter of 8-mm full width at half maximum.

The time points in each voxel were detrended for correcting the linear drift over time. Nuisance signals, including whole brain, white matter, cerebrospinal fluid and six motion parameters, were regressed out from rs-fMRI time courses [27,28]. Subsequently, temporal filtering with a band of 0.01-0.08 Hz was applied to the time courses to reduce the impact of low-frequency drifts and high-frequency noise [29]. Then, the above pre-processed rs-fMRI data were used for FC analysis.

### Region of interest-based FC analysis

Stemming from the well-defined neuroanatomy of DMN [30,31], five pairs of brain regions are selected as regions of interest (ROIs) and then generated by the WFU Pick Atlas toolbox [32] (Fig. 1a) (http://fmri.wfubmc.edu/software/PickAtlas): including medial prefrontal cortex (mPFC), lateral temporal cortex (LTC), posterior cingulate cortex (PCC), inferior parietal lobe (IPL), hippocampal formation (HF). To better control the heterogeneity in cortical anatomy across individuals, we used a relatively large seed (10 mm) in order to elicit a broader connectivity map.

**Figure. 1.**
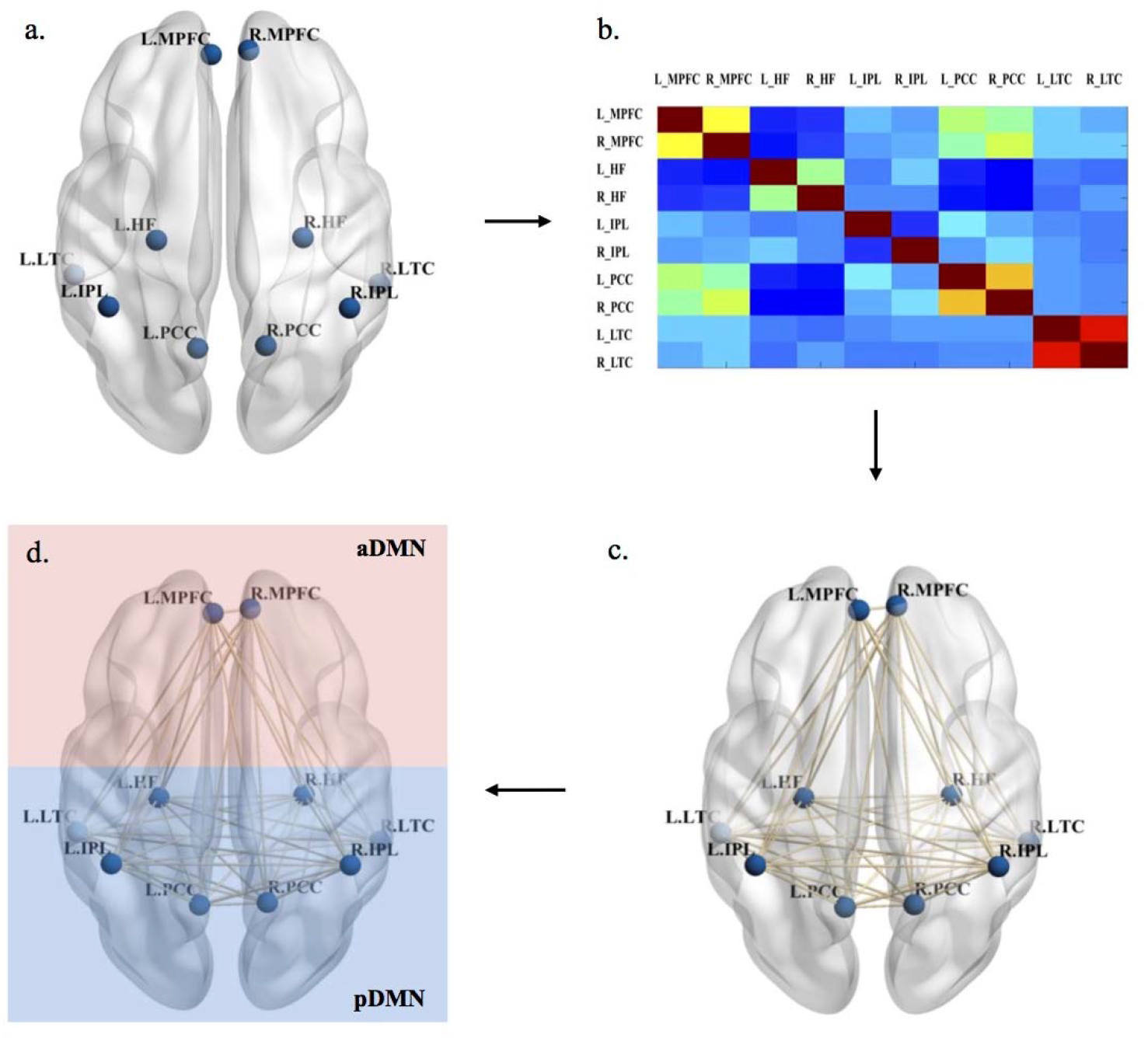
The procedures employed for mapping the connectivity within default mode network (DMN). including (a) defining the regions of interest (ROIs) of DMN regions; (b) constructing individual correlation matrix; (c) calculating the functional connectivity strength (FCS) within DMN and (d) demonstrating the anterior and posterior DMN. The visualization of ROIs and connections were performed by the Brain Net Viewer (Available at: https://www.nitrc.org/projects/bnv/).

After defining the ROIs, the mean time courses were extracted from each ROI by averaging the time courses over all voxels within the ROI. To illustrate the organization within DMN, Pearson correlation coefficients were calculated with the corrected time courses between the ROIs (Fig. 1b). The absolute values of the correlation coefficient (*r*) were used as a measure to evaluate the FCS between the ROIs (Fig. 1c) [33–35]. Smaller value of FCS indicates the reduced FCS within DMN.

### Statistical analyse

Comparisons of demographics and scores of neurocognitive tasks across groups were tested either with *χ^2^* for categorical variable or ANOVA for continuous variables. When appropriate, the Tukey method was conducted to do the *post hoc* multiple comparisons. Effect size was evaluating using the Partial eta squared (*η*^2^). As to ROI-wise FC analysis, the correlation matrix of ROIs was performed individually with family false discovery rate (FDR) correction. General linear model (GLM) was conducted to test the main effects and interaction effects of APOE *ε* 4 and cognitive status. Pearson correlation coefficient was used to assess the relationship between FCS and cognitive performance. ANOVA, *χ^2^* test and Pearson correlation coefficients were performed by SPSS Statistics (Version 20, IBM. Armonk NY, USA). All tests were two-tailed, and the significance level threshold was set at *p* < 0.05 unless explicitly stated otherwise.

## RESULTS

### Demographics and cognitive performance

According to APOE *ε* 4 genotype and cognitive status, the participants were divided into four groups: CN adults without APOE *ε* 4 (CN *ε* 4-), CN adults with APOE *ε* 4 (CN *ε* 4+), MCI patients without APOE *ε* 4 (MCI *ε* 4-) and MCI patients with APOE *ε* 4 (MCI *ε* 4+). A shown in Table 1, the demographics in terms of age (*F* = 2.434, *p* = 0.073) and sex (*F* = 0.768, *p* = 0.516) were comparable between groups. Compared with CN adults, MCI patients had fewer years of education and worse performance on global cognitive efficiency (ADAS-Cog: *F* = 21.726, *p* < 0.001), working memory function (Delayed recall of words: *F* = 42.057, *p* < 0.001) and executive function (CVFT: *F* = 4.575, *p* = 0.006).

**Table 1.** Demographics and cognitive characteristics across groups

| | CN $\varepsilon$ 4-<br>(n=27) | CN $\varepsilon$ 4+<br>(n=11) | MCI $\varepsilon$ 4-<br>(n=17) | MCI $\varepsilon$ 4+<br>(n=10) | *F* | *P value* |
| --- | --- | --- | --- | --- | --- | --- |
| Age | 70.39±4.35 | 69.48±4.41 | 72.67±3.52 | 72.92±2.99 | 2.434 | 0.073 |
| Gender (M/F) | 10/17 | 4/7 | 8/6 | 6/4 | 0.768 | 0.516 |
| Education (years) | 8.48±3.87 | 7.01±4.26 | 7.17±4.71 | 4.21±4.16 | 2.801 | 0.047 |
| CMMSE | 27.81±2.38 | 27.36±3.01 | 25.58±3.18 | 25.61±2.59 | 3.115 | 0.033 |
| ADAS-Cog | 6.18±2.85 | 7.36±2.24 | 13.58±4.12 | 11.11±2.76 | 21.726 | <0.001 |
| Delayed recall | 6.67±1.52 | 6.09±1.31 | 2.12±1.49 | 2.71±1.56 | 42.057 | <0.001 |
| DSF | 4.07±1.84 | 3.64±1.29 | 2.81±1.21 | 2.51±1.27 | 2.841 | 0.045 |
| DSB | 3.63±1.55 | 3.45±1.75 | 3.73±1.75 | 5.31±1.25 | 3.707 | 0.016 |
| CVFT | 39.89±6.41 | 41.36±10.54 | 32.88±10.61 | 31.91±6.55 | 4.575 | 0.006 |
| TMT-A (second) | 15.71±20.43 | 13.27±7.87 | 18.44±11.04 | 21.41±15.93 | 0.543 | 0.655 |
| TMT-B (second) | 20.56±14.01 | 17.27±10.64 | 27.06±25.55 | 28.61±16.17 | 1.196 | 0.319 |

Note. Data are raw scores and presented as mean ± SD.
CMMSE = Cantonese mini mental state examination; ADAS-Cog = Alzheimer's Disease Assessment Scale cognitive subscale; DSF = Digit span forward; DSB = Digit span backward; CVFT = Category verbal fluency test; TMT = Trail making test.

### Results of volumetric analysis

With a threshold of uncorrected *p* < 0.001, there was no voxel left. Altthough APOE carriers had less grey matter volume than non-carriers (CN *ε* 4-: 579.85 ± 39.19, CN *ε* 4+: 574.17 ± 42.98, MCI *ε* 4-: 557.91 ± 65.08, MCI *ε* 4+: 544.99 ± 38.06), there was no statistically significant difference across four groups (*F* = 1.139, *p* = 0.341).

### Functional connectivity within DMN

Among 45 pairwise FC within DMN (Supplementary Table), significant group-wise differences were found in the FCS of left HF and right HF (*F* = 4.735, *p* = 0.005, corrected), right HF and left LTC (*F* = 3.589, *p* = 0.019, corrected), left PCC and right PCC (*F* = 3.435, *p* = 0.023, corrected), left PCC and left LTC (*F* = 3.657, *p* = 0.017, corrected) (Table 2). Post hoc analysis showed that the FCS between left HF and right HF, and the FCS between left PCC and right PCC were reduced in MCI *ε* 4+ group. In contrast, MCI *ε* 4+ group showed increased FCS between left PCC and left LTC than the CN *ε* 4+ group. Compared with non-carriers and CN adults, APOE *ε* 4 carriers with MCI showed reduced connectivity within DMN, particularly the interhemispheric ones.

**Table 2.** Functional connectivity strength within DMN across groups

| | CN $\varepsilon$ 4-<br>(n=27) | CN $\varepsilon$ 4+<br>(n=11) | MCI $\varepsilon$ 4-<br>(n=17) | MCI $\varepsilon$ 4+<br>(n=10) | *F* | *P value* |
| --- | --- | --- | --- | --- | --- | --- |
| L.HF and R.HF | 1.18±0.44 | 0.95±0.56 | 0.79±0.35 | 0.69±0.46 | 4.735 | 0.005 |
| R.HF and L.LTC | 0.58±0.34 | 0.36±0.21 | 0.32±0.17 | 0.51±0.33 | 3.589 | 0.019 |
| L.PCC and R.PCC | 1.57±0.25 | 1.33±0.38 | 1.44±0.31 | 1.21±0.51 | 3.435 | 0.023 |
| L.PCC and L.LTC | 0.29±0.23 | 0.16±0.14 | 0.19±0.14 | 0.43±0.28 | 3.657 | 0.017 |

Note. Data are raw scores and presented as mean ± SD.
Abbreviations: HF = Hippocampal formation; LTC = Lateral temporal cortex; PCC = Posterior cingulate cortex.

### Effect of APOE ε 4 and cognitive status on FCS

As Fig. 2 shown, using age, sex and years of education as covariates, the main effect of APOE *ε* 4 was found in the FC of left PCC and right PCC (*F* = 6.133, *p* = 0.016, *η*^2^ = 0.096); the main effect of cognitive status was found in the FC of right mPFC and left IPL (*F* = 4.585, *p* = 0.036, *η*^2^ = 0.073). The interaction effect between APOE *ε* 4 and cognitive status was found in the FC of right mPFC and left LTC (*F* = 4.698, *p* = 0.034, *η*^2^ = 0.075), right HF and left LTC (*F* = 7.673, *p* = 0.008, *η*^2^ = 0.117), left PCC and left LTC (*F* = 6.719, *p* = 0.012, *η*^2^ = 0.104), right IPL and right LTC (*F* = 4.516, *p* = 0.038, *η*^2^ = 0.072).

**Figure. 2.**
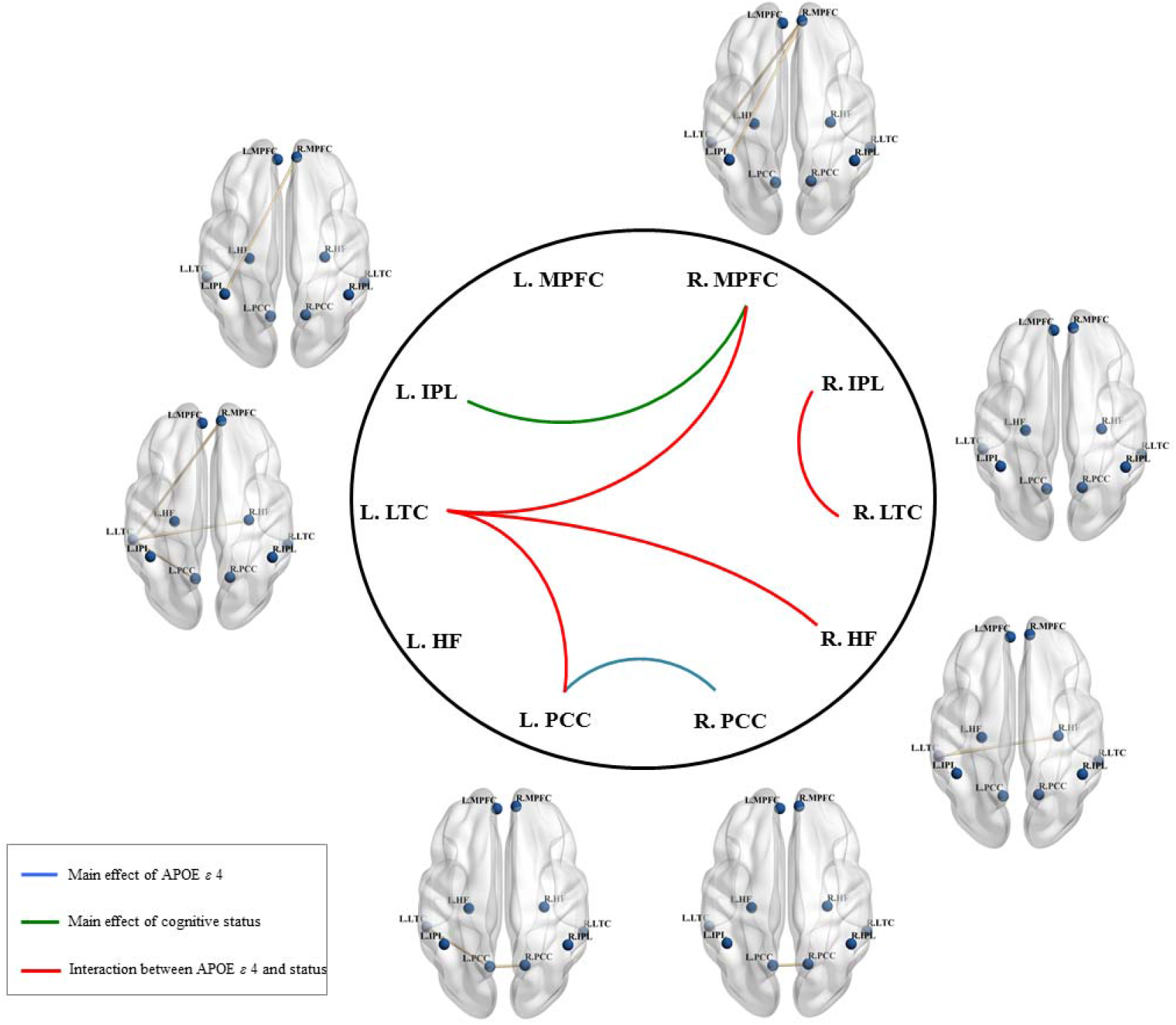
Main effect and interaction effect of APOE *ε* 4 and cognitive status on the functional connectivity strength (FCS) within DMN. Green line represents the main effect of cognitive status; blue line represents the main effect of APOE *ε* 4; red line represents interaction effect. Abbreviations: mPFC = Medial prefrontal cortex; IPL = inferior parietal lobe; HF = Hippocampal formation; LTC = Lateral temporal cortex; PCC = Posterior cingulate cortex.

### Association between FCS and cognitive performance

The FCS of left mPFC and right IPL (*r* = 0.303, *p* = 0.019, corrected) and the FCS of left mPFC and right LTC (*r* = 0.316, *p* = 0.015, corrected) were correlated with better CVFT performance (Fig. 3a). The FCS of left IPL and right IPL (*r* = 0.326, *p* = 0.012, corrected) and the FCS of left IPL and right PCC (*r* = 0.285, *p* = 0.029, corrected) were correlated with better performance of delayed recall (Fig. 3b).

**Figure. 3.**
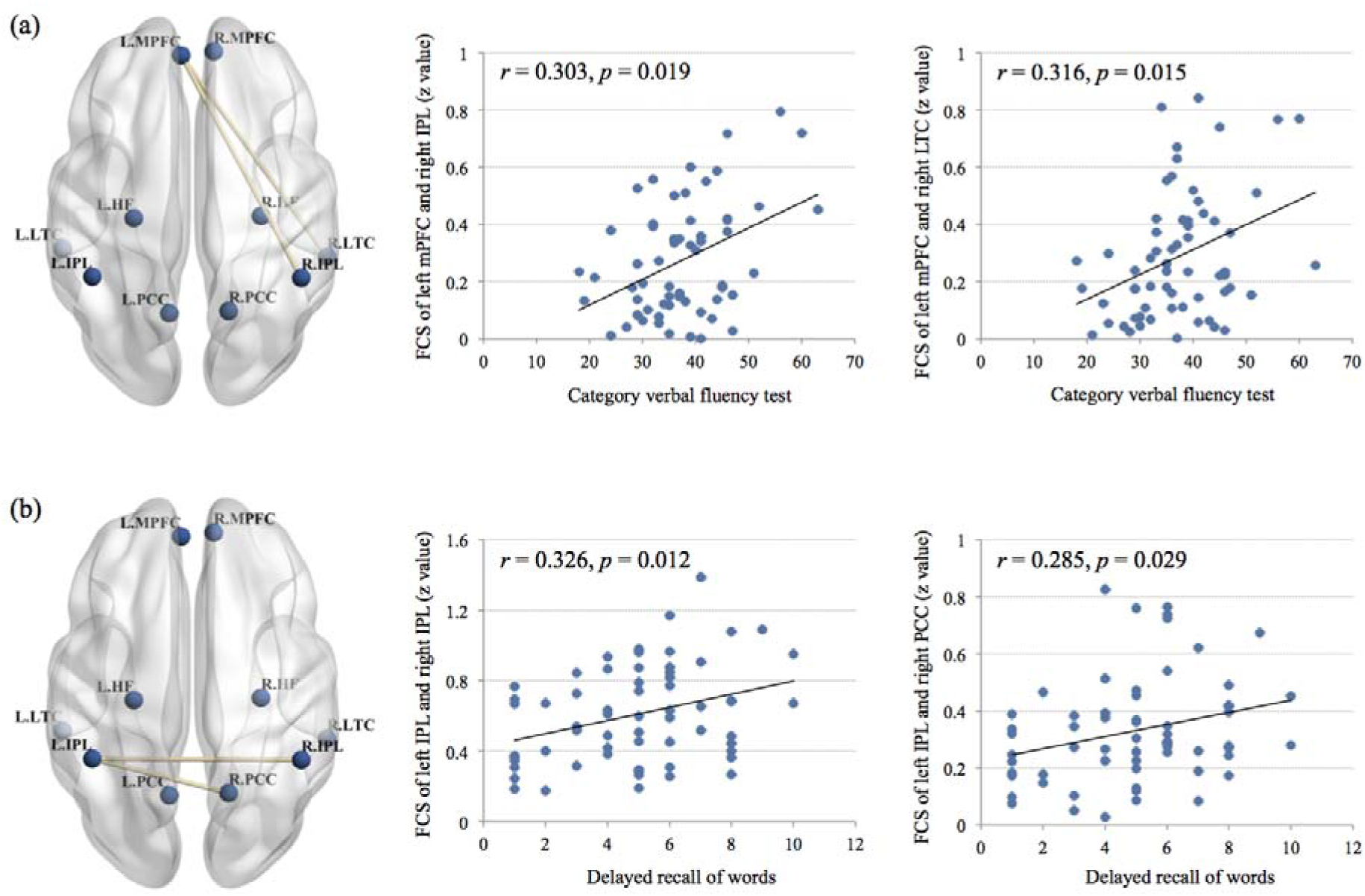
Associations between functional connectivity strength (FCS) and cognitive functions. (a) The FCS of left mPFC and right IPL and the FCS of left mPFC and right LTC were correlated with better performance of CVFT; (b) the FCS of left IPL and right IPL and the FCS of left IPL and right PCC were correlated with better performance of delayed recall. Abbreviations: mPFC = Medial prefrontal cortex; IPL = inferior parietal lobe; HF = Hippocampal formation; LTC = Lateral temporal cortex; PCC = Posterior cingulate cortex.

## DISCUSSION

We identified the main effects of APOE *ε* 4 and cognitive status and their interaction effects on the connectivity within DMN with functional connectivity strength (FCS) and then examined its associations with cognitive performance in senior individuals with or without genetic risk for AD. Overall, APOE *ε* 4 MCI carriers had disturbed functional organization within DMN than CN adults and MCI non-carriers. Cognitive status-related effect was found on aDMN (i.e., FC between left IPL and right mPFC). APOE *ε* 4-related effect and the interaction effects of APOE *ε* 4 and cognitive status were found mainly on pDMN (i.e., FC between left PCC and right PCC).

Several mechanisms might contribute to our findings of differential effect within DMN. First, age, APOE *ε* 4 and cognitive status may selectively affect the DMN connectivity. For instance, network-level alternations of DMN connectivity might be due to the accumulations of cognitive dysfunction, including age-related [36], aging-related [4] and AD-related cognitive decline [37]. Specifically, the abnormal activation in left hippocampus, prefrontal and parietal regions during memory-related task in APOE *ε* 4 carriers [38] has been confirm by rs-fMRI studies with functional connectivity mapping [39]. In line with most previous findings [17,21], we observed the effect of genetic risks (i.e., APOE *ε* 4) was on the connectivity between bilateral PCC and cognitive status was related to the reduced connectivity of frontoparietal areas, of which the cortical areas are vulnerable to AD pathology [40]. Another possibility is that the disturbed connectivity may reflect the domain-specific brain functions in late adulthood. For example, Harrison et al observed reduced connectivity within pDMN in older adults during memory encoding and this detectable disturbance was associated with poor cognitive performance [41]. In a longitudinal fucntional investigation, changes in anterior-posterior DMN connectivity were specifically associated with episodic memory [4].

Beyond the main effects of APOE *ε* 4 and cognitive status on DMN connectivity, another results deserved to highlight here are the interaction effects between these two factors on brain connectivity. There was a specific pattern of the interaction effects on DMN connectivity driven almost by the interhemisphric FC linked with left temporal cortex, of which has no overlapping with either APOE *ε* 4-related or cognitive status-related FC. Interestingly, lateral temporal cortex plays a key role in the process of working memory [42] and language [43], of which the function is highly lateralized [44]. Indeed, the strong associations to memory (i.e., delayed recall), language (i.e., CVFT) and DMN connectivity changes of the regions identified as significantly different between groups in this study converge on the potential importance of these regions and the effect of APOE *ε* 4 on their cognitive function.

In conclusion, functional connectivity strength, as a quantifiable connectivity measure related to network-level brain connections, can help us to better understand the complex relationship between age, genetic risk factor and brain function. The status-specific patterns the interaction effects on the connectivity within DMN can be used as the early neurophenotypes of AD progression, which may highlight its potential use in further develping network-guided non-invasive brain stimulation and other advanced brain interventions for cognitive enhancement.

### Limilations and future directions

Some limitations of this study need to be acknowledged: (1) the small sample size of APOE *ε* 4 MCI carriers is the main weakness of this study; (2) the backgrounds of cerebrovascular risk factors are not included, which might affect the functional connectivity as well; (3) we only focused on the genetic risk of APOE *ε* 4 allele and DMN; hence the potential effects of APOE *ε* 2 on the other brain network were not launched.

There is still much to learn about the disease-specific intrinsic brain network changes in types of neurodegenerative disorders and about how brain connectivity and neuropathology in relation to each other over time. We would use these changes to detect the individuals at risk of cognitive deficits and further guide the personalized intervention, such as non-invasive brain stimulation.

## DISCLOSURE OF INTEREST

The authors declare that they have no competing interest.

## ACKNOWLEDGEMENTS

This research was supported by the Direct Grant from the Chinese University of Hong Kong. The authors would like to express their gratitude to Prof. Yong He, Dr. Mingrui Xia, Dr. Zaixu Cui, Dr. Zhengjia Dai and Dr. Xuhong Liao from the State Key Laboratory of Cognitive Neuroscience and Learning & IDG/McGovern Institute for Brain Research, Beijing Normal University for their great suggestion of functional connectivity analysis. Additionally, the authors also thank all the subjects who are willing to participate in our study for their indispensable contribution to the field of research.

---

**Supplementary Table.** The 45 pairs of functional connectivity strength within DMN

| | CN $\varepsilon$ 4-<br>(n=27) | CN $\varepsilon$ 4+<br>(n=11) | MCI $\varepsilon$ 4-<br>(n=17) | MCI $\varepsilon$ 4-+<br>(n=10) | *F* | *P value* |
| --- | --- | --- | --- | --- | --- | --- |
| L.mPFC_R.mPFC | 1.26±0.31 | 1.35±0.41 | 1.49±0.35 | 1.20±0.46 | 1.68 | 0.18 |
| L.mPFC_L.HF | 0.22±0.20 | 0.23±0.20 | 0.23±0.18 | 0.25±0.12 | 0.07 | 0.98 |
| L.mPFC_R.HF | 0.17±0.18 | 0.24±0.19 | 0.25±0.18 | 0.35±0.28 | 1.89 | 0.14 |
| L.mPFC_L.IPL | 0.30±0.23 | 0.37±0.29 | 0.31±0.20 | 0.36±0.28 | 0.36 | 0.78 |
| L.mPFC_R.IPL | 0.27±0.19 | 0.37±0.30 | 0.24±0.18 | 0.29±0.19 | 0.93 | 0.43 |
| L.mPFC_L.PCC | 0.42±0.27 | 0.48±0.31 | 0.36±0.17 | 0.34±0.29 | 0.63 | 0.60 |
| L.mPFC_R.PCC | 0.31±0.21 | 0.39±0.29 | 0.36±0.24 | 0.24±0.17 | 0.91 | 0.44 |
| L.mPFC_L.LTC | 0.28±0.21 | 0.21±0.19 | 0.25±0.14 | 0.33±0.30 | 0.59 | 0.63 |
| L.mPFC_R.LTC | 0.27±0.23 | 0.38±0.30 | 0.27±0.13 | 0.29±0.31 | 0.56 | 0.64 |
| R.mPFC_L.HF | 0.24±0.18 | 0.29±0.20 | 0.25±0.17 | 0.36±0.16 | 1.14 | 0.34 |
| R.mPFC_R.HF | 0.22±0.16 | 0.18±0.15 | 0.21±0.18 | 0.32±0.28 | 1.19 | 0.32 |
| R.mPFC_L.IPL | 0.32±0.24 | 0.36±0.21 | 0.34±0.19 | 0.49±0.26 | 1.40 | 0.25 |
| R.mPFC_R.IPL | 0.31±0.26 | 0.38±0.26 | 0.20±0.10 | 0.29±0.18 | 1.46 | 0.24 |
| R.mPFC_L.PCC | 0.43±0.26 | 0.44±0.32 | 0.39±0.19 | 0.31±0.29 | 0.65 | 0.58 |
| R.mPFC_R.PCC | 0.38±0.22 | 0.43±0.32 | 0.36±0.18 | 0.28±0.19 | 0.80 | 0.50 |
| R.mPFC_L.LTC | 0.37±0.23 | 0.34±0.24 | 0.21±0.23 | 0.48±0.44 | 2.15 | 0.10 |
| R.mPFC_R.LTC | 0.24±0.20 | 0.46±0.32 | 0.29±0.19 | 0.31±0.31 | 2.16 | 0.10 |
| L.HF_R.HF | 1.18±0.44 | 0.95±0.57 | 0.72±0.31 | 0.69±0.46 | 4.74 | 0.005 |
| L.HF_L.IPL | 0.37±0.27 | 0.25±0.20 | 0.38±0.27 | 0.35±0.28 | 0.65 | 0.59 |
| L.HF_R.IPL | 0.32±0.22 | 0.29±0.38 | 0.40±0.27 | 0.40±0.28 | 0.59 | 0.62 |
| L.HF__L.PCC | 0.32±0.28 | 0.23±0.17 | 0.23±0.28 | 0.25±0.14 | 0.66 | 0.58 |
| L.HF__R.PCC | 0.28±0.24 | 0.23±0.19 | 0.26±0.22 | 0.24±0.21 | 0.13 | 0.94 |
| L.HF__L.LTC | 0.50±0.39 | 0.54±0.27 | 0.31±0.19 | 0.39±0.31 | 1.49 | 0.23 |
| L.HF__R.LTC | 0.44±0.26 | 0.47±0.33 | 0.31±0.37 | 0.37±0.21 | 0.86 | 0.47 |
| R.HF__L.IPL | 0.39±0.28 | 0.17±0.11 | 0.43±0.34 | 0.46±0.39 | 2.23 | 0.09 |
| R.HF__R.IPL | 0.37±0.24 | 0.27±0.24 | 0.39±0.27 | 0.34±0.30 | 0.51 | 0.68 |
| R.HF__L.PCC | 0.28±0.22 | 0.26±0.22 | 0.17±0.14 | 0.28±0.20 | 1.01 | 0.39 |
| R.HF__R.PCC | 0.26±0.24 | 0.27±0.20 | 0.15±0.13 | 0.25±0.22 | 0.96 | 0.42 |
| R.HF__L.LTC | 0.58±0.34 | 0.36±0.21 | 0.31±0.17 | 0.51±0.33 | 3.40 | 0.019 |
| R.HF__R.LTC | 0.47±0.34 | 0.40±0.32 | 0.48±0.35 | 0.52±0.26 | 0.25 | 0.86 |
| L.IPL__R.IPL | 0.70±0.31 | 0.53±0.29 | 0.56±0.22 | 0.63±0.24 | 1.39 | 0.26 |
| L.IPL__L.PCC | 0.33±0.18 | 0.37±0.28 | 0.25±0.11 | 0.29±0.26 | 0.84 | 0.48 |
| L.IPL__R.PCC | 0.33±0.18 | 0.38±0.26 | 0.21±0.14 | 0.33±0.23 | 1.67 | 0.18 |
| L.IPL__L.LTC | 0.38±0.27 | 0.33±0.22 | 0.28±0.24 | 0.36±0.25 | 0.48 | 0.70 |
| L.IPL__R.LTC | 0.37±0.29 | 0.20±0.13 | 0.29±0.23 | 0.40±0.30 | 1.49 | 0.23 |
| R.IPL__L.PCC | 0.34±0.25 | 0.39±0.27 | 0.22±0.19 | 0.28±0.30 | 1.22 | 0.31 |
| R.IPL__R.PCC | 0.37±0.29 | 0.45±0.32 | 0.26±0.17 | 0.31±0.27 | 1.15 | 0.34 |
| R.IPL__L.LTC | 0.47±0.33 | 0.35±0.30 | 0.21±0.20 | 0.33±0.26 | 2.68 | 0.06 |
| R.IPL__R.LTC | 0.35±0.22 | 0.48±0.24 | 0.42±0.31 | 0.27±0.19 | 1.68 | 0.18 |
| L.PCC__R.PCC | 1.57±0.25 | 1.33±0.38 | 1.39±0.29 | 1.20±0.51 | 3.44 | 0.02 |
| L.PCC__L.LTC | 0.29±0.23 | 0.16±0.14 | 0.19±0.14 | 0.43±0.28 | 3.66 | 0.02 |
| L.PCC__R.LTC | 0.22±0.19 | 0.22±0.22 | 0.26±0.20 | 0.22±0.13 | 0.12 | 0.95 |
| R.PCC__L.LTC | 0.27±0.22 | 0.24±0.24 | 0.22±0.19 | 0.31±0.24 | 0.37 | 0.78 |
| R.PCC__R.LTC | 0.25±0.22 | 0.31±0.30 | 0.23±0.21 | 0.16±0.11 | 0.80 | 0.50 |
| L.LTC__R.LTC | 0.62±0.40 | 0.66±0.65 | 0.48±0.29 | 0.66±0.50 | 0.48 | 0.70 |

Abbreviations: mPFC = Medial prefrontal cortex; IPL = inferior parietal lobe; HF = Hippocampal formation; LTC = Lateral temporal cortex; PCC = Posterior cingulate cortex.

